# *Hras^G12V^* induces follicular thyroid cancer with attenuated MAPK activation and increased latency compared to *Kras^G12D^*

**DOI:** 10.64898/2026.09.09.750413

**Authors:** Nicholas E. Bambach, Katherine Labella, Lee Ann Jolly, Diana Isabel Cruz, Nicole Massol, Antonio Di Cristofano, Aime T. Franco

## Abstract

RAS mutations are found in nearly 50% of follicular thyroid cancers (FTCs), frequently accompanied by secondary mutations in the PI3K/AKT pathway as tumors advance to more poorly differentiated states. To examine the role that oncogenic *Hras* plays in thyroid tumor initiation and progression, we developed murine models with thyroid-specific expression of *Hras^G12V^* combined with heterozygous or homozygous loss of *Pten.* Loss of *Pten* cooperated with *Hras^G12V^* in a dose-dependent manner to induce the development of follicular thyroid carcinoma and poorly-differentiated thyroid carcinoma. Histopathology of *Hras^G12V^/Pten^Hom^* tumors closely resembled those from the established *Kras^G12D^/Pten^Hom^* model, but tumor onset was significantly delayed in *Hras^G12V^/Pten^Hom^* mice. At three weeks of age, downregulation of MAPK pathway inhibitors was observed in *Kras^G12D^/Pten^Hom^* thyroids, accompanied by increased MAPK pathway activation compared to *Hras^G12V^/Pten^Hom^* mice. Furthermore, amplification of oncogenic *Ras* was found in *Hras^G12V^* tumors and cell lines, while allelic balance was maintained in *Kras^G12D^* models. These studies demonstrate clear phenotypic differences between mutant *Hras* and *Kras* in the thyroid and suggest that delayed MAPK activation may mediate the increased tumor latency observed in the *Hras^G12V^/Pten^Hom^* model.

## Introduction

Over the last four decades, the prevalence of thyroid cancer has steadily increased, solidifying it as the most common endocrine malignancy (1,2). Presently, thyroid cancer is predicted to become the fourth most frequently diagnosed cancer in the United States by 2030 (3). More than 95% of patients are diagnosed with well-differentiated tumors such as papillary thyroid cancer (PTC) or follicular thyroid cancer (FTC) (4). However, a small subset of patients progresses to poorly differentiated thyroid cancer (PDTC) or anaplastic thyroid cancer (ATC), which together are responsible for the majority of thyroid cancer-related mortality (5,6). Standard treatment for thyroid cancer is surgical resection and often radioactive iodine therapy, leading to the necessity of lifetime thyroid hormone replacement. Although tumor-associated mortality of thyroid cancer is low, morbidity is high. This underscores the need for a more thorough understanding of the role of different oncogenes and cooperative mutations in mediating disease pathogenesis to develop more precise treatment strategies that will improve patients’ survivorship and long-term quality of life.

In the last decade, the field of knowledge surrounding the genetic alterations associated with thyroid cancer has expanded dramatically. Activation of the mitogen-activated protein kinase (MAPK) signaling pathway is frequently observed across the spectrum of thyroid tumors, from benign adenomas to advanced anaplastic disease. Patients often present with somatic gain-of-function mutations in *BRAF* or *RAS* family genes (7–10). *BRAF^V600E^* is the most common alteration found in PTC, whereas activating mutations in *HRAS*, *KRAS*, and *NRAS* are particularly common both in follicular adenomas and follicular carcinomas (reviewed in (11)). Though *KRAS* alterations are the most frequently observed in other cancer types, mutations in *HRAS* and *NRAS* are predominant in the thyroid (7–10). While the prevalence of specific *RAS* isoform mutations has been well characterized, the distinct role of these isoforms in the thyroid remains unexplored. Under normal circumstances, RAS family members are ubiquitously expressed but their activation is tightly regulated. However, mutations in RAS can lead to production of a constitutively active form of the protein, activating the MAPK pathway. While this activation is known to drive thyroid tumor initiation and maintenance, the mechanisms by which individual RAS driver mutations impact thyroid tumorigenesis remains underexplored.

Much like MAPK, activation of the phosphoinositide 3-kinase (PI3K)-signaling cascade is common in thyroid cancer, especially in advanced disease (12–14). There is considerable evidence, both from basic studies as well as human genetic data, that aberrant PI3K-AKT signaling may cooperate with MAPK activation and mediate the progression from well-differentiated to advanced carcinoma (15–17). Notably, *PTEN* pathogenic variants, such as those observed in *PTEN* hamartoma tumor syndrome (PHTS), can predispose patients to thyroid tumors (18). Therefore, loss of *PTEN* provides an alternative and clinically relevant mechanism to activate the PI3K pathway and can be used in model systems of thyroid disease. Indeed, previous studies demonstrated that activation of the PI3K pathway through loss of *Pten* cooperates with MAPK signaling to rapidly induce tumorigenesis in *Braf^V600E^*-driven tumor models (19,20). Furthermore, Miller, et. al., showed that *Pten* loss cooperates with expression of *Kras^G12D^* to induce murine thyroid tumors (20). While *KRAS^G12D^* mutations predominate in most cancers, they are relatively rare in thyroid tumors (21). Therefore, we aimed to evaluate whether thyroid-specific expression of oncogenic *Hras* combined with heterozygous or homozygous loss of *Pten* would cooperate in the development of FTC in a manner similar to what has been observed with oncogenic *Kras*.

Here, we report that *Hras^G12V^* cooperates with *Pten* loss to induce the development of murine FTCs and PDTCs. We observed that *Hras^G12V^* transforms thyroid tissue with equal severity and penetrance as previously described in *Kras^G12D^* models (20). However, the latency of disease progression was much greater with *Hras^G12V^*-mutant tumors. Unlike their *Kras^G12D^* counterparts, early-timepoint *Hras^G12V^* thyroids display upregulation of ERK inhibitors such as *Dusp* family members. Consistent with these findings, we observed earlier MAPK activation in *Kras^G12D^*-driven lesions compared to *Hras^G12V^*. Tumor-derived *Kras^G12D^/Pten^Hom^* cell lines similarly displayed increased MAPK activation and were more resistant to MEK inhibition. Additionally, *Hras^G12V^*-mutant tumors displayed an allelic imbalance not observed in their wildtype or *Kras^G12D^*-mutant counterparts, suggesting that Hras-mediated transformation may depend on allele dosage. Consistent with this, human molecular data from the TCGA showed that *HRAS* expression correlated with a decreased levels of thyroid differentiation and increased levels of MAPK activation, commonly associated with more aggressive disease and worsened response to treatment (22,23). Our phenotypic and molecular findings underscore differences in the oncogenic capacity of HRAS and KRAS, and offer insights into how these results may translate to patients.

## Materials and Methods

### Experimental animals and hormone measurements

*FR-Hras^G12V^*, *Pten^flox/flox^*, and thyroid peroxidase promoter (TPO)-Cre mice have been previously described (24–26). Mice originated on a mixed genetic background and were subsequently backcrossed into the 129/SvImJ (Jackson Laboratory mouse strain #002448) background for at least 9 generations to create 129Sv-*Hras^G12D^/Pten^Het^* and 129Sv-*Hras^G12D^/Pten^Hom^* lines.

*Kras^G12D^/Pten^Hom^* mice have been previously described (20) and were maintained on a 129Sv background. Genotypes were determined by PCR using previously described primers (20,24). All procedures were approved by the Institutional Animal Care Committee of the University of Arkansas for Medical Sciences or Albert Einstein School of Medicine.

Blood from mice was collected via cardiac puncture immediately after euthanasia with CO2 and centrifuged at maximum speed at 4°C for 15 minutes. Serum was removed and stored at -80°C until assayed. Serum TSH levels were determined as previously described (27). The lower limit of detection for TSH in this assay was 10 mU/l. The serum levels of total thyroxine were measured using an antibody-coated tube radioimmunoassay (RIA) (Siemens Medical Solutions Diagnostics) adapted for mouse serum. The lower limit of detection for levothyroxine (L-T4) in this assay was 0.25 μg/dL.

### Histology and immunohistochemistry

Thyroid tissue was fixed in 10% buffered formalin and embedded in paraffin. 5 μm-thick sections were prepared and stained with hemotoxylin and eosin. Histological diagnosis was performed by a thyroid pathologist (NM) blinded to the genotype. Paraffin-embedded tissues were dewaxed in xylene and rehydrated in alcohol. Following citrate antigen retrieval (Thermo Scientific), tissue sections were blocked with goat serum and incubated overnight at 4°C with the following antibodies at the specified dilutions: Ki-67 (#9129, 1:200), phospho-p44/42 MAPK (#9101, 1:200), and phospho-Akt S473 (#9271, 1:200) (all from Cell Signaling Technology). Staining with diaminobenzidine (DAB) was performed using the Elite ABC kit (Vector Labs) according to the manufacturer’s protocol and visualized on an Olympus DP73 microscope equipped with a Nikon Eclipse 8400 camera. Images were acquired using Cell Sens Entry software (Olympus).

### Cell culture, growth curves, and thyrosphere assays

Cell lines were established from thyroid tumors from *Kras^G12D^/Pten^Hom^* (T683, T826) (28) and 129Sv*-Hras^G12V^/Pten^Hom^* (HRAS1, H340T) double-mutant mice and grown in F12 medium in the presence of 10% fetal bovine serum (FBS). All lines were confirmed to be of thyroid origin and devoid of stromal contamination by verifying complete recombination of *Pten* via PCR. Primer sequences are listed in the Supplementary Materials and Methods. Cells for growth curves were plated (500 cells/well in 10% FBS) in 96-well plates. At the indicated time points, nuclei were stained with Hoechst 33342 (Thermo Scientific) diluted at 1:200 in culture medium. The stained nuclei were imaged with the EVOS M7000 and cell counts were quantified using ImageJ. Counts were averaged across 10 replicate wells per cell line. Cells for thyrospehere assays were seeded (1000 cells/well in 10% FBS) in quadruplicate in 24-well ultra-low attachment plates (Corning). Cells were cultured for seven days and spheroids were imaged with the EVOS M7000. Spheroid count and size were quantified using Fiji.

### RNA isolation, RT-qPCR, and RT-qPCR arrays

Thyroid lobes were surgicallyremoved and immediately placed in liquid nitrogen. RNA was isolated using TRIzol Reagent (Thermo Fisher Scientific) and the Direct-zol RNA Miniprep Kit (Zymo Research). 1000 ng of RNA was reverse transcribed using the Verso cDNA Synthesis Kit (Thermo Scientific) in the presence of random hexamers. Reverse transcription quantitative PCR (RT-qPCR) reactions were performed using PowerTrack SYBR Green Master Mix (Applied Biosystems), the Quant Studio Real-Time PCR System (Applied Biosystems), and primer pairs for *Ccnd1*, *Mapk12*, *Dusp1*, *Dusp3*, *Dusp4*, and *Dusp6*. *Actb* was used to normalize the expression of all samples. Primer sequences are listed in the Supplementary Materials and Methods.

The RT^2^ Profiler Mouse MAPK PCR Array (Qiagen) was used to analyze the expression of 84 MAPK pathway-related genes in *Kras^G12D^/ Pten^Hom^* and *Hras^G12V^/Pten^Hom^* cell lines and wildtype thyroids. The PrimePCR Array (Bio-Rad) was used to analyze the expression of 40 genes related to inhibition of the ERK pathway in thyroids from 3-week-old wildtype, 3-week-old *Kras^G12D^/ Pten^Hom^,* 3-week-old *Hras^G12V^/Pten^Hom^*, and 1-year-old *Hras^G12V^/Pten^Hom^* animals. Briefly, total RNA from cell lines and mouse thyroids was extracted and purified using TRIzol Reagent and the RNeasy Plus Mini Kit (Qiagen). RNA was reverse transcribed with the RT^2^ First Strand Kit (Qiagen) or Verso cDNA Synthesis Kit (Thermo Scientific), combined with SYBR Green Master Mix (Applied Biosystems), and added to each well of the array plate containing predispensed gene-specific primer sets. Expression was determined using the Ct method and normalized using four independent housekeeping genes. Results were validated on additional independent samples by RT-qPCR as described above.

### Western blot analysis

Cells were lysed in radioimmunoprecipitation assay (RIPA) buffer supplemented with 1X Halt Protease and Phosphatase Inhibitor Cocktail and phenylmethylsulfonyl fluoride (PMSF) (Thermo Scientific). Total protein was quantified via bicinchoninic acid (BCA) assay, and 10 µg of protein was separated by SDS-PAGE. Protein was transferred to a PVDF membrane (Millipore) and blocked with 5% BSA for 1 hour at room temperature. Membranes were incubated with the primary antibody overnight at 4°C, secondary antibody for 1 hour at room temperature, and visualized using SuperSignal West Femto Maximum Sensitivity Substrate (Thermo Scientific).

The following antibodies were purchased from Cell Signaling Technology and used at the following concentrations: GAPDH (#2118, 1:5000), AKT (#9272, 1:2000), phospho-AKT S473 (#9271, 1:2000), phospho-AKT T308 (#9275, 1:2000), p44/42 MAPK (#9102, 1:2000), phospho-p44/42 MAPK (#9101, 1:2000), p38 gamma MAPK (#2307, 1:2000), Cyclin D1 (#2922, 1:2000), HRP-linked Goat anti-Rabbit IgG (#7074, 1:5000).

### Dose-response assays

HRAS1, H340T, T683, and T826 cells (1000/well in 10% FBS) were seeded in 96-well plates.

After 24 hours, cells were dosed with the indicated concentrations of LY294002 (Selleck Chemicals) or selumetinib (Selleck Chemicals). Cells were cultured for 72 hours and nuclei were stained with Hoechst 33342 (Thermo Scientific) diluted at 1:200 in culture medium. The stained nuclei were then imaged with the EVOS M7000 and cell counts were quantified using ImageJ.

All experiments were performed with six replicates per concentration. Cell counts were normalized and the IC50 for each cell line was calculated using GraphPad’s four-parameter logistic regression.

### Allele-specific polymerase chain reaction

Thyroid lobes and tail clippings were taken from mice at the time of sacrifice and placed on ice. Genomic DNA was isolated using the Quick-DNA Miniprep Kit (Zymo Research) according to the manufacturer’s instructions. Polymerase chain reaction (PCR) was performed to amplify both the mutant and wildtype alleles of *Hras* and *Kras.* Primer sequences are provided in the Supplementary Materials and Methods. PCR products were resolved by agarose gel electrophoresis and imaged using the iBright CL1500 Imaging System (Invitrogen). The presence of mutant and wild-type alleles was determined based on the expected amplicon sizes. Gels were analyzed using Fiji’s gel analyzer densitometry function and the ratio of mutant to wild-type band intensity was calculated for each lane.

### Thyroid differentiation, ERK, and BRAF-RAS scores

RNA expression data from samples in the TCGA-THCA cohort was accessed and downloaded through the cBioPortal (29). Thyroid differentiation (TDS), enhanced thyroid differentiation (eTDS), ERK, and BRAF-RAS scores were calculated as previously described (30,31).

Associations between *HRAS* and *KRAS* mRNA expression and molecular scores were assessed using Pearson correlation analysis.

### Statistical analysis

All statistical analyses were conducted using GraphPad Prism 10 (v10.4.1) or RStudio (v4.4.2).

Fisher’s exact test, Pearson correlation, one-way ANOVA, and log-rank testing were performed as appropriate. A two-sided *p* value < 0.05 was considered statistically significant.

## Results

### *Pten* dosage dictates malignant progression in *Hras^G12V^*-driven thyroid carcinoma

To investigate how PI3K pathway activation cooperates with constitutively active Hras to promote thyroid cancer initiation and progression, we used an *in vivo* approach that closely recapitulates the expression of oncogenic *Hras^G12V^* and loss of *Pten* in patients. We crossed *FR-Hras^G12V^/Pten^flox/flox^* mice with *Pten^flox/+^/TPO-Cre* mice, which express Cre recombinase under the control of the human thyroid peroxidase (TPO) promoter, to generate a thyrocyte-specific heterozygous knock-in of oncogenic *Hras^G12V^* with concurrent heterozygous (*Hras*^G12V^/*Pten^Het^*) or homozygous (*Hras*^G12V^/*Pten^Hom^*) deletion of *Pten* (Figure 1A)*. Hras^G12V^/Pten^Het^* and *Hras^G12V^/Pten^Hom^* mice were born at the expected Mendelian frequency, and we observed no differences in weight or outward appearance between wild-type and any transgenic animals (Figure 1B). While previous studies have found that activation of the MAPK pathway can dysregulate thyroid hormone production (20,32–34), we observed no significant differences in serum LT4 or TSH levels between control and *Hras^G12V^/Pten^Het^* or *Hras^G12V^/Pten^Hom^* mice (Figure S1). These results support our prior observations that endogenous expression of *Hras^G12V^* does not alter expression of genes involved with thyroid hormone biosynthesis or responsiveness to TSH (32). Consistent with normal thyroid function, *Hras^G12V^/Pten^Het^* and *Hras^G12V^/Pten^Hom^* animals were fertile and produced viable, healthy litters which were successfully weaned.

**Figure 1.**
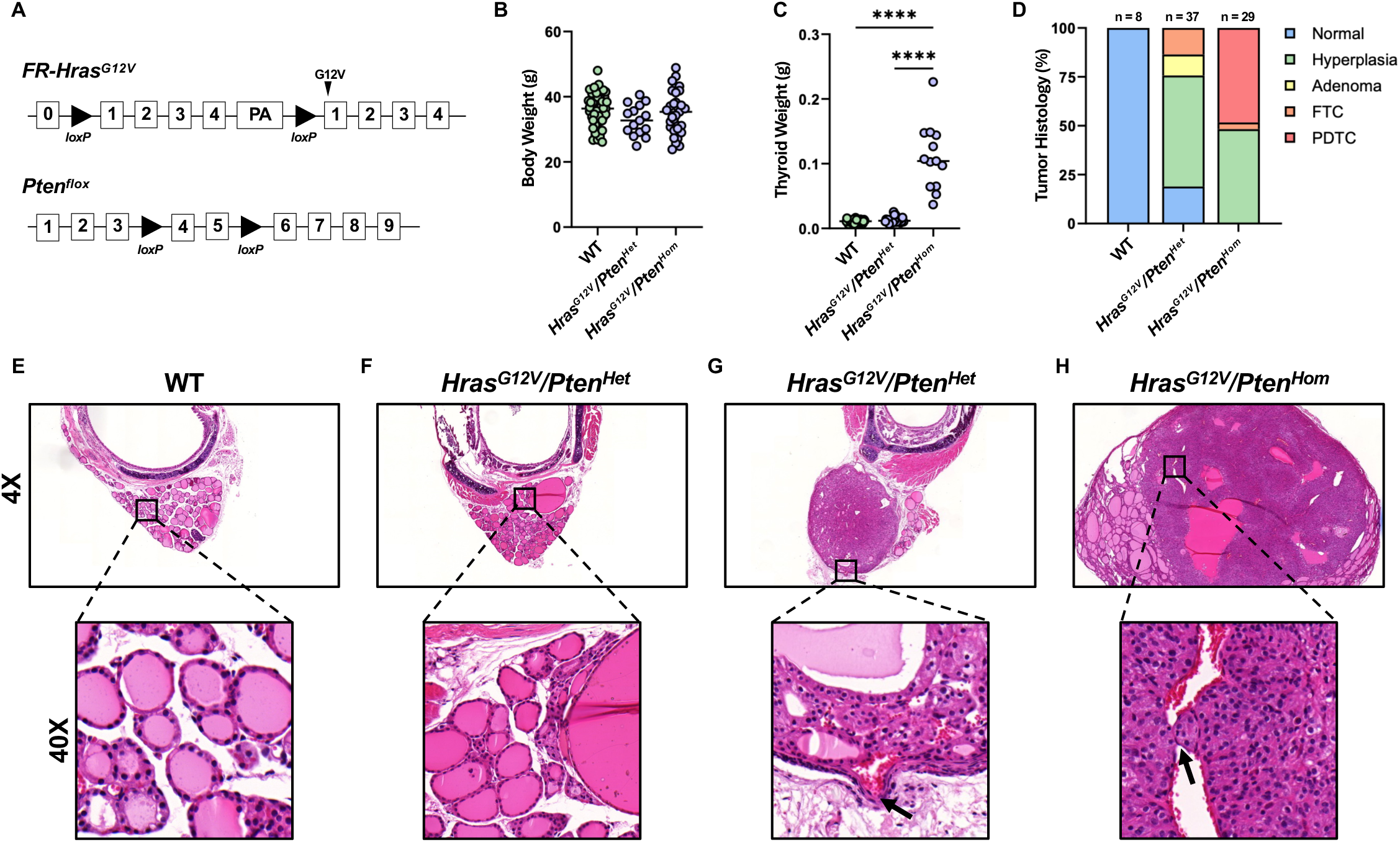
**(A)** Schematic of *Hras^G12V^* and *Pten^flox^* genetic constructs utilized to generate experimental model. **(B)** Body weights of 1-year-old wild-type (n = 51), *Hras^G12V^/Pten^Het^* (n = 15), and *Hras^G12V^/Pten^Hom^* (n = 31) animals. **(C)** Thyroid weights of 1-year-old wild-type (n = 40), *Hras^G12V^/Pten^Het^* (n = 27), and *Hras^G12V^/Pten^Hom^* (n = 13) animals. **(D)** Histological diagnosis of extracted thyroids from wild-type, *Hras^G12V^/Pten^Het^*, and *Hras^G12V^/Pten^Hom^* animals. Diagnosis was performed by a thyroid pathologist (NM) blinded to the genotype. **(E-H)** Representative images at low (4X) and high (40X) magnification of H&E-stained thyroids extracted from experimental animals. **(E)** Normal thyroid containing follicles filled with colloid from a wild-type mouse. **(F)** Benign hyperplasia in a thyroid from an *Hras^G12V^/Pten^Het^* mouse. **(G)** Follicular thyroid carcinoma (FTC) with capsular invasion (arrow) in a thyroid from an *Hras^G12V^/Pten^Het^* mouse. **(H)** Poorly differentiated thyroid carcinoma (PDTC) with vascular invasion (arrow) in a thyroid from an *Hras^G12V^/Pten^Hom^* mouse. One-way ANOVA **** *p* < 0.0001.

To assess the impact of *Pten* dosage on tumor formation, we first compared the weights of thyroids from 1-year-old wild-type, *Hras^G12V^/Pten^Het^*, and *Hras^G12V^/Pten^Hom^* animals. *Hras^G12V^/Pten^Hom^* thyroids were significantly larger than wild-type and *Hras^G12V^/Pten^Het^* thyroids (Figure 1C), suggesting pathologic enlargement of the gland. Extracted thyroids were then stained with hematoxylin and eosin for histological analysis. All wild-type animals retained normal thyroid histology and exhibited no evidence of pathologic changes (Figure 1D, E). *Hras^G12V^/Pten^Het^* mice developed predominantly benign lesions, with 56.8% exhibiting thyroid hyperplasia and 10.8% forming benign adenomas (Figure 1D,F). Benign tumors were characterized by nodular hyperplasia with Hürthle cell adenomas and oncocytic changes. Five animals (13.5%) developed well-differentiated follicular thyroid cancer (FTC), identified by classical features such as capsular invasion (Figure 1D,G). At one year, the latest time-point evaluated, seven animals (18.9%) remained disease free with normal thyroid histology. (Figure 1D). In contrast, *Hras^G12V^/Pten^Hom^* mice had higher incidence of malignant transformation (OR = 6.63, 95% CI = 1.85-28.15, *p* = 0.0011). Approximately 50% of mice developed poorly differentiated thyroid carcinoma (PDTC) (Figure 1D, H). Tumors were characterized by numerous mitoses, atypical mitoses, nuclear pleomorphism, necrosis, and vascular invasion (Figure 1H, Figure S2). All *Hras^G12V^/Pten^Hom^* animal displayed evidence of abnormal thyroid histology, supporting that homozygous loss of *Pten* enhances thyroid transformation (Figure 1D).

### *Hras^G12V^*-driven tumors exhibit increased latency compared to *Kras^G12D^*-driven tumors

We next sought to determine whether our *Hras^G12V^/Pten^Hom^*-driven model phenotypically recapitulated the previously established *Kras^G12D^/Pten^Hom^*-driven model of FTC (20). Interestingly, while *Kras^G12D^/Pten^Hom^* and *Hras^G12V^/Pten^Hom^* mice develop tumors that are histopathologically similar, the penetrance of disease significantly differed. *Kras^G12D^/Pten^Hom^* mice were reported to develop tumors with 100% penetrance and complete lethality by 20 weeks of age (20). In contrast, only 51.7% of *Hras^G12V^/Pten^Hom^* animals developed malignant disease by one year of age and exhibited no early lethality (Figure 1D, Table 1). We hypothesized that the observed differences between the two models may be attributable to modifiers present in their distinct genetic backgrounds. Whereas *Kras^G12D^/Pten^Hom^* mice were maintained on a congenic 129Sv background, *Hras^G12V^/Pten^Hom^* animals were generated on a mixed genetic background. Therefore, *Hras^G12V^/Pten^Hom^* animals were backcrossed onto the 129Sv background for nine generations. SNP analysis confirmed that 129Sv-*Hras^G12V^/Pten^Het^* and 129Sv-*Hras^G12V^/Pten^Hom^* animals were over 99% identical to the parental 129Sv background.

**Table 1.** Summary of malignant tumor formation and lung metastases in mice of different ages.

| Strain | Age (Weeks) | Malignant Tumor | Lung Metastasis |
| --- | --- | --- | --- |
| *Hras<sup>G12V</sup>/Pten<sup>Hom</sup>* | 20 | 0% (0/9) | 0% (0/9) |
|  | 52 | 51.7% (15/29) | 0% (0/29) |
| 129Sv-*Hras<sup>G12V</sup>/Pten<sup>Hom</sup>* | 20 | 0% (0/10) | 0% (0/10) |
|  | 52 | 100% (13/13) | 76.9% (10/13) |
| *Kras<sup>G12D</sup>/Pten<sup>Hom</sup>* (12) | 20 | 100% | 100% |

Interestingly, after backcrossing, the penetrance of disease increased in 129Sv-*Hras^G12V^/Pten^Hom^* mice whereby 100% of mice developed malignant tumors by one year of age (Table 1). The resultant tumors from congenic 129Sv-*Hras^G12V^/Pten^Hom^* animals were histologically similar to those developed on the mixed background. While the penetrance of the backrossed *Hras^G12V^*-driven model was now identical to that of the *Kras^G12D^*-driven model, 129Sv-*Hras^G12V^/Pten^Hom^* mice exhibited increased disease latency. *Kras^G12D^/Pten^Hom^* mice were reported to rapidly develop FTC, which progressed to PDTC, and resulted in complete mortality by 20 weeks (20). In contrast, 20-week-old 129Sv-*Hras^G12V^/Pten^Hom^* mice displayed no overt phenotype and no evidence of pathologic transformation (Table 1). In addition to harboring advanced thyroid tumors, all *Kras^G12D^/Pten^Hom^* animals were reported to develop lung metastases (20). We therefore evaluated the presence of metastatic disease in *Hras^G12V^/Pten^Hom^* animals. Consistent with the lack of primary tumor formation, 20-week-old *Hras^G12V^*-mutant animals exhibited no evidence of lung metastases (Table 1). However, by 52 weeks, 76.9% of *Hras^G12V^/Pten^Hom^* mice harbored lung macrometastases (Table 1, Figure S3). The increased latency of tumor and metastatic lesion formation in *Hras^G12V^*-mutant animals compared to *Kras^G12D^*-mutant animals, despite a similar penetrance and identical genetic background, suggests that oncogenic *Kras* drives tumorigenesis more rapidly than *Hras*.

### *Kras^G12D^* enhances MAPK signaling and MEK inhibitor resistance compared to *Hras^G12V^*

To further investigate the mechanisms underlying the observed phenotypic differences, stable cell lines were derived from *Hras^G12V^/Pten^Hom^* tumors (HRAS1 and H340T) and *Kras^G12D^/Pten^Hom^* tumors (T683 and T826 (28)). The cell lines were confirmed to be of thyroid lineage and devoid of stromal cell contamination by verifying complete *Pten* recombination (Figure S4A). Cell lines were evaluated for the presence of cancer stem cells or self-renewing populations by assessing their ability to form thyrospheres in culture (35). Both *Kras^G12D^-* and *Hras^G12V^*-driven cell lines formed similar numbers of thyrospeheres when plated on low-adherence tissue culture plates (Figure S4B). However, *Kras^G12D^* spheroids were significantly larger than *Hras^G12V^* spheroids, suggesting that despite having the same capacity for self-renewal, *Kras^G12D^*-driven cells may proliferate faster than *Hras^G12V^*-driven ones, possibly contributing to the accelerated tumor formation observed in *Kras^G12D^/Pten^Hom^* animals (Figure 2A, B). We next examined the proliferation of these cell lines and found that despite the differences in spheroid size, *Hras^G12V^*- and *Kras^G12D^-*driven cells grew at similar rates in 2D culture (Figure S4C).

**Figure 2.**
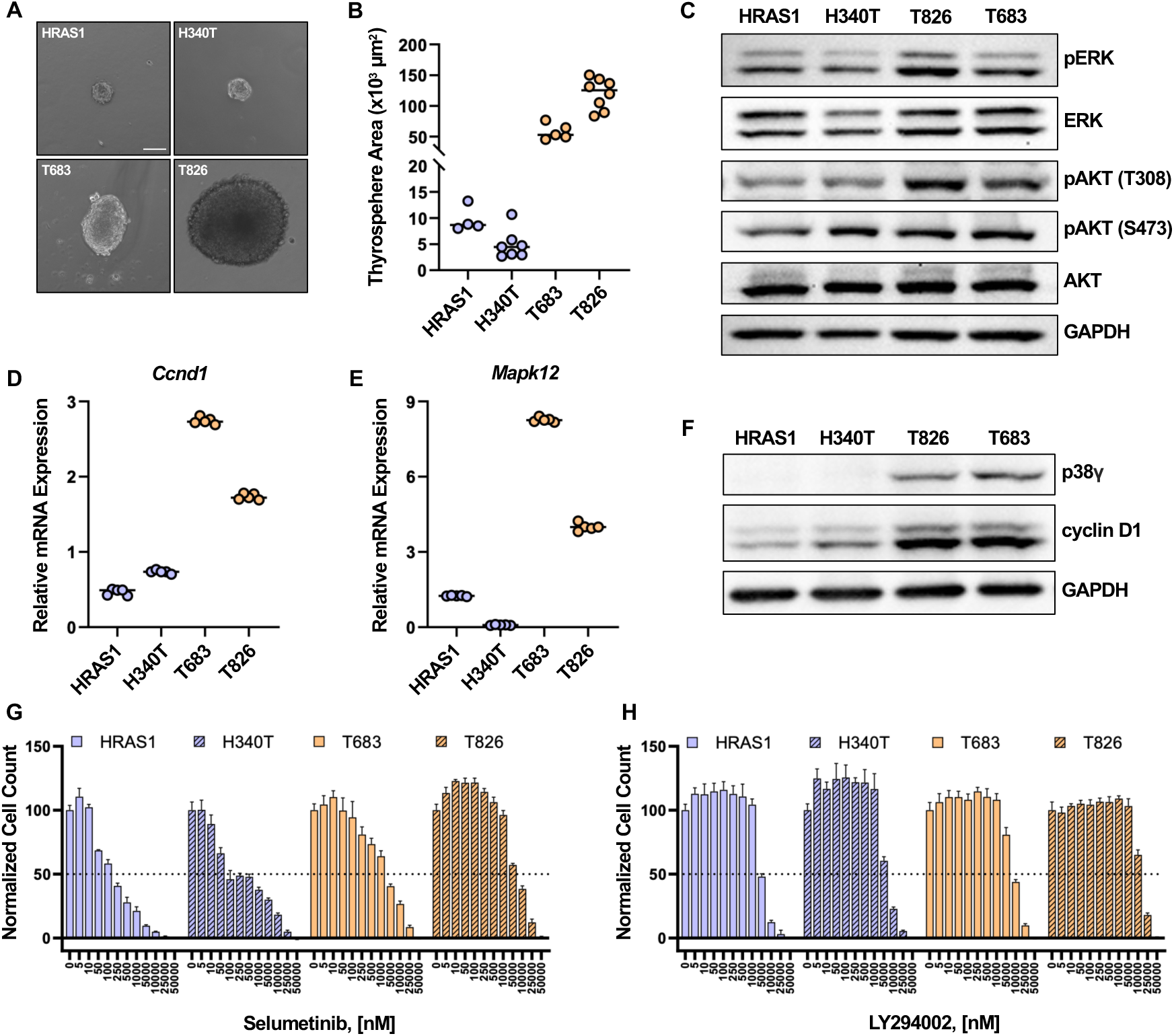
**(A)** Representative microscopy images of thyrospheres formed from HRAS1, H340T, T683, and T826 cell lines after seven days in culture in low-attachment conditions. Scale bar = 100 µm. **(B)** Area of thyrospheres formed from HRAS1 (n = 4), H340T (n = 7), T683 (n = 5), and T826 (n = 8). **(C)** Western blot analysis of phosphorylated ERK and AKT in HRAS1, H340T, T683, and T826 cell lines. **(D-E)** RT-qPCR analysis of *Ccnd1* and *Mapk12* expression in HRAS1, H340T, T683, and T826 cell lines. **(F)** Western blot analysis of cyclin D1 (*Ccnd1*) and p38γ *(Mapk12)* in HRAS1, H340T, T683, and T826 cell lines. **(G)** Viability of HRAS1, H340T, T683, and T826 cells treated with varying concentrations of the MEK inhibitor selumetinib. **(H)** Viability of HRAS1, H340T, T683, and T826 cells treated with varying concentrations of the PI3K inhibitor LY294002. Dose-response experiments were performed with six replicates per concentration.

To determine whether oncogenic *Hras^G12V^* and *Kras^G12D^* induce differential activation of mitogenic signaling pathways, we evaluated the phosphorylation levels of immediate MAPK and PI3K downstream effectors. We found that both *Kras^G12D^-*driven cell lines had increased phosphorylation of ERK and the Thr308 residue of AKT compared to the *Hras-*driven HRAS1 and H340T (Figure 2C). Interestingly, T826 had more robust activation of ERK and AKT than T683, suggesting underlying heterogeneity even in the presence of the same genetic driver. To further profile the molecular changes associated with *Kras^G12D^* and *Hras^G12V^*, an RT-qPCR array was utilized to measure the expression levels of 84 genes associated with MAPK signaling in T683 and HRAS1 cell lines. Each cell line had a distinct expression pattern that was unique compared to wild-type thyroids (Figure S4D). T683 exhibited increased expression of multiple genes involved in cell cycle progression, including cyclins *(Ccnd1, Ccne1, Ccna2, Ccnb1, Ccnb2)* and cyclin dependent kinases *(Cdk2, Cdk4)* compared to HRAS1. Additionally, T683 exhibited upregulation of several MAPK family members, including those involved in the p38 MAPK pathway *(Mapk11, Mapk12, Mapk14, Map2k3, Mapkap2)*. To validate these findings, RT-qPCR and Western blotting analysis of two selected genes were performed across all four cell lines. We found that cyclin D1 *(Ccnd1)* and p38γ *(Mapk12)* were upregulated in both *Kras-* driven cell lines compared to the *Hras-*driven ones (Figure 2D-F). Together, these results indicate that *Kras^G12D^* induces more robust activation of the MAPK pathway than *Hras^G12V^ in vitro*.

Due to the differences in basal MAPK and PI3K signaling levels, we investigated whether *Hras*- and *Kras*-driven cell lines differ in their sensitivity to MAPK and PI3K pathway inhibition. We tested the efficacy of selumetinib and LY294002, two potent inhibitors of MEK and PI3K. Selumetinib inhibited HRAS1 and H340T at IC50 values of 170.6 nM and 279.3 nM, respectively (Figure 2G). In contrast, both *Kras*-driven cell lines exhibited resistance to MEK inhibition, with IC50 values of 2,246 nM and 6,763 nM for T683 and T826, respectively (Figure 2G). Notably, the greater resistance of T826 compared to T683 is consistent with the higher basal levels of ERK and AKT phosphorylation observed by Western blot analysis. A moderate increase in the IC50 of LY294002 was also observed in *Kras-*driven cell lines compared to *Hras*-driven lines. LY294002 inhibited HRAS1 and H340T with IC50 values of 4.92 µM and 6.09 µM, respectively, whereas the *Kras*-driven cell lines T683 and T826 exhibited higher IC50 values of 9.13 µM and 13.32 µM, respectively (Figure 2H).

### *Kras^G12D^* induces more rapid MAPK activation than *Hras^G12V^* during tumorigenesis

Having shown that *Kras^G12D^* drives stronger mitogenic signaling than *Hras^G12V^* in fully transformed cells, we tested whether similar differences are present during early tumorigenesis, which might account for the delayed tumor onset in *Hras^G12V^*-mutant animals.

Immunohistochemical staining of downstream MAPK and PI3K effectors was performed in thyroid tissue from 3-week-old *Hras^G12V^/Pten^Hom^* and *Kras^G12V^/Pten^Hom^* mice. Levels of pERK and pAKT were significantly higher in *Kras^G12D^/Pten^Hom^* thyroid tissue than in *Hras^G12V^/Pten^Hom^* tissue (Figure 3A). Consistent with increased levels of mitogenic signaling, cells in *Kras^G12D^/Pten^Hom^* thyroids were also significantly more proliferative, as measured by increased Ki67 positivity (Figure 3A, B). These findings suggest that *Kras^G12D^* more rapidly promotes stronger MAPK and PI3K signaling and a more proliferative state, which may contribute to the accelerated transformation observed in *Kras*-mutant animals.

**Figure 3.**
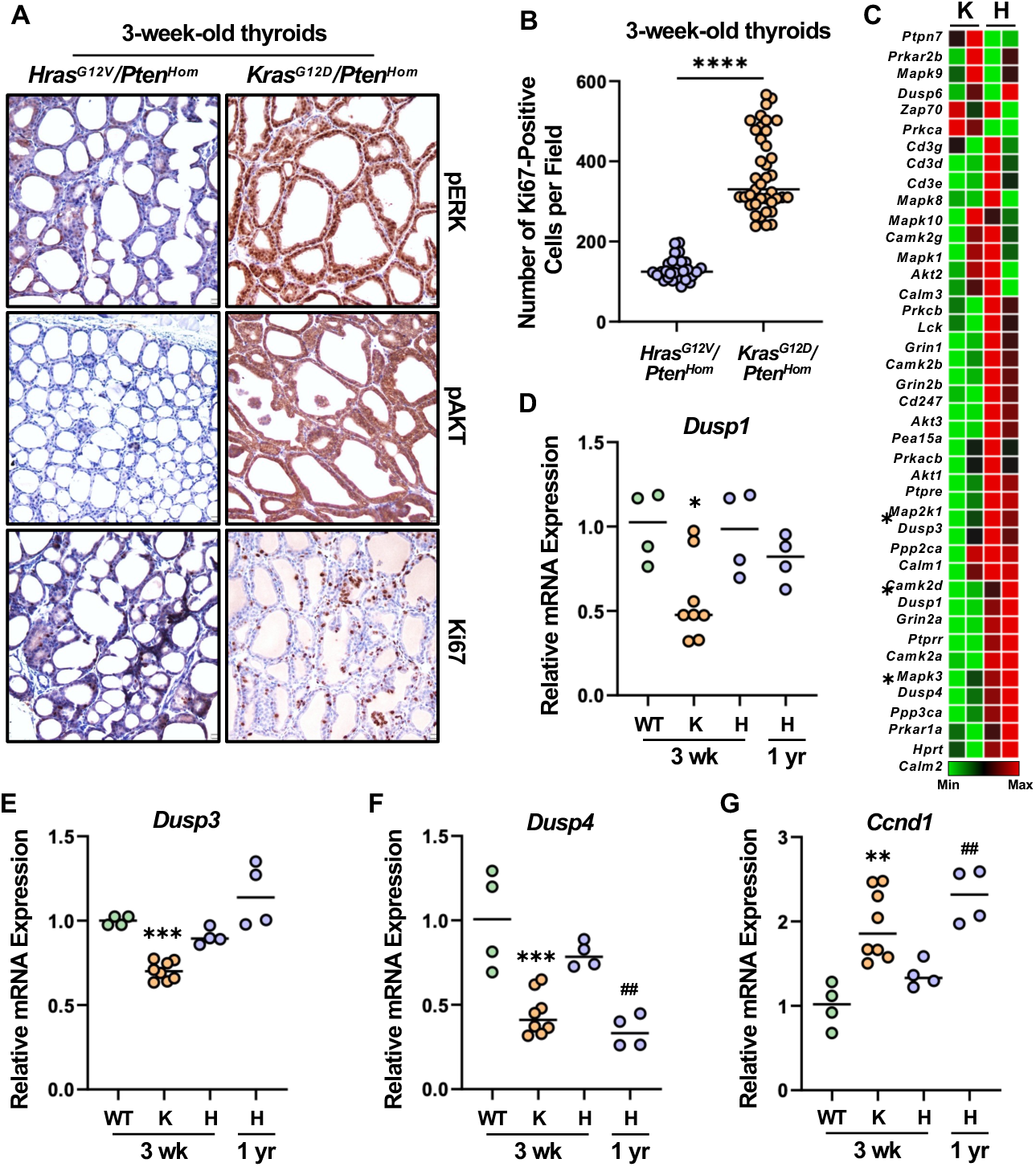
**(A)** Representative immunohistochemical staining of pERK, pAKT, and Ki67 in thyroid tissue from 3-week-old *Hras^G12V^/Pten^Hom^* and *Kras^G12D^/Pten^Hom^* mice. **(B)** Quantification of Ki67-positive cells per 10X field in thyroid tissue from 3-week-old *Hras^G12V^/Pten^Hom^* and *Kras^G12D^/Pten^Hom^* animals. Student’s t-test **** *p* < 0.0001. **(C)** Expression of MAPK regulatory genes in thyroid tissue from 3-week-old *Kras^G12D^/Pten^Hom^* (K) and *Hras^G12V^/Pten^Hom^* (H) mice. Expression was measured using a 40 gene PCR array with two replicates per genotype. Genes further validated via RT-qPCR are denoted by asterisks. **(D-G)** RT-qPCR analysis of *Dusp1, Dusp3, Dusp4,* and *Ccnd1* expression in thyroid tissue from 3-week-old wild-type (n = 4), *Kras^G12D^/Pten^Hom^* (n = 8), and *Hras^G12V^/Pten^Hom^* (n = 4) mice and 1-year-old *Hras^G12V^/Pten^Hom^* (n = 4) mice. One-way ANOVA * *p* < 0.05, ** *p* < 0.01, *** *p* < 0.001 versus wild-type; ## *p* < 0.01 versus 3-week-old *Hras^G12V^/Pten^Hom^*.

We next investigated whether the observed differences in MAPK activation were associated with altered expression of genes involved in regulation of the MAPK pathway. We performed a PCR array to assess the expression of MAPK regulatory genes in thyroids from 3-week-old *Hras^G12V^/Pten^Hom^* and *Kras^G12V^/Pten^Hom^* mice. Several genes encoding proteins that inhibit the MAPK pathway were downregulated in 3-week-old *Kras^G12D^/Pten^Hom^* thyroids compared to *Hras^G12V^/Pten^Hom^* thyroids, including the phosphatases *Dusp1*, *Dusp3*, and *Dusp4*, as well as *Ptprr*, *Ppp2ca*, and *Pea15a* (Figure 3C). Among these, members of the dual-specificity phosphatases (DUSP) family were of particular interest as they attenuate signaling by directly dephosphorylating activated MAPKs. In a normal state, DUSPs are induced by MAPK signaling and function as a negative feedback mechanism to restrain pathway activity; however, this feedback loop is frequently uncoupled in cancer, resulting in impaired DUSP expression and sustained MAPK signaling (36–41).

To validate the array, RT-qPCR analysis of *Dusp1*, *Dusp3*, and *Dusp4* expression was performed using additional independent thyroid samples from 3-week-old wildtype, *Kras^G12D^/Pten^Hom^* and *Hras^G12V^/Pten^Hom^* mice. *Dusp1, Dusp3,* and *Dusp4* expression were significantly decreased in *Kras^G12D^/Pten^Hom^* thyroids compared to both wildtype and *Hras^G12V^/Pten^Hom^* tissue, suggesting loss of the canonical negative feedback loop, potentially contributing to the elevated MAPK signaling induced by *Kras^G12D^* (Figure 3D-F). In contrast, expression of *Dusp1, Dusp3,* and *Dusp4* in 3-week-old *Hras^G12V^/Pten^Hom^* thyroid tissue was comparable to that of wild-type thyroids, indicating an intact feedback mechanism in *Hras^G12V^*-mutant cells (Figure 3D-F). To determine whether *Dusp* expression changes in *Hras^G12V^*-driven tumors at the time of complete transformation, we analyzed thyroid tissue from 1-year-old *Hras^G12V^/Pten^Hom^* mice. *Dusp4* expression was diminished compared to 3-week-old *Hras^G12V^*-mutant thyroids, consistent with reduced MAPK-dependent feedback regulation during tumor progression (Figure 3F).

Coincident with reduced *Dusp4* levels, expression of the MAPK-responsive gene *Ccnd1* was also increased in 1-year-old *Hras^G12V^/Pten^Hom^* thyroids compared to 3-week-old animals, reaching levels comparable to those observed in 3-week-old *Kras^G12D^ /Pten^Hom^* thyroids (Figure 3G). Together, these results identify marked temporal differences in MAPK pathway activity and feedback regulation between *Kras^G12D^* and *Hras^G12V^* after oncogene activation.

### Allelic amplification is specific to *Hras^G12V^* and not *Kras^G12D^* during thyroid transformation

We have previously reported that homozygous knock-in of *Hras^G12V^* alone is insufficient to transform thyrocytes (32), and that amplification of the mutant *Hras^G12V^* allele is required to induce skin papilloma formation (24,42). We therefore hypothesized that amplification of *Hras^G12V^* occurs in the context of homozygous *Pten* loss to drive malignant transformation. To determine whether amplification of *Hras^G12V^* is present at the time of complete transformation, we examined the *Hras* allelic ratio in tumors from 1-year-old *Hras^G12V^/Pten^Hom^* mice. The mutant *Hras^G12V^* allele was amplified in thyroid tumors (Figure 4A, C). In contrast, paired tail DNA exhibited the expected approximately 1:1 ratio of wild-type and targeted alleles (Figure 4A, C). Since we observed this imbalance at the time of complete transformation, we next examined the allelic ratio in thyroids prior to transformation. In 3-week-old thyroids from *Hras^G12V^/Pten^Hom^* mice, the wild-type and mutant *Hras* alleles were present at the expected approximately 1:1 ratio in both thyroid tissue and paired tail DNA, suggesting that *Hras^G12V^* amplification occurs during the course of malignant transformation (Figure 4B,C). To then directly test whether increased *Hras^G12V^* dosage drives accelerated tumorigenesis, we generated mice with homozygous knock-in of oncogenic *Hras^G12V^* (*Hras^G12V/G12V^/Pten^Hom^*). In stark contrast to mice heterozygous for *Hras^G12V^*, all *Hras^G1V2/G12V^/Pten^Hom^* animals rapidly developed tumors and died by 40 weeks of age (Figure 4D). To assess whether this allelic amplification was specific to *Hras^G12V^,* we analyzed tumors from *Kras^G12D^/Pten^Hom^* mice and found that the ratio of wild-type to mutant *Kras* was equivalent (Figure S5). These results suggest that *Kras^G12D^* is sufficient to drive rapid thyroid transformation without undergoing amplification. In contrast, *Hras^G12V^* may require increased allele dosage to fully induce transformation, possibly contributing to the increased latency in tumor development.

**Figure 4.**
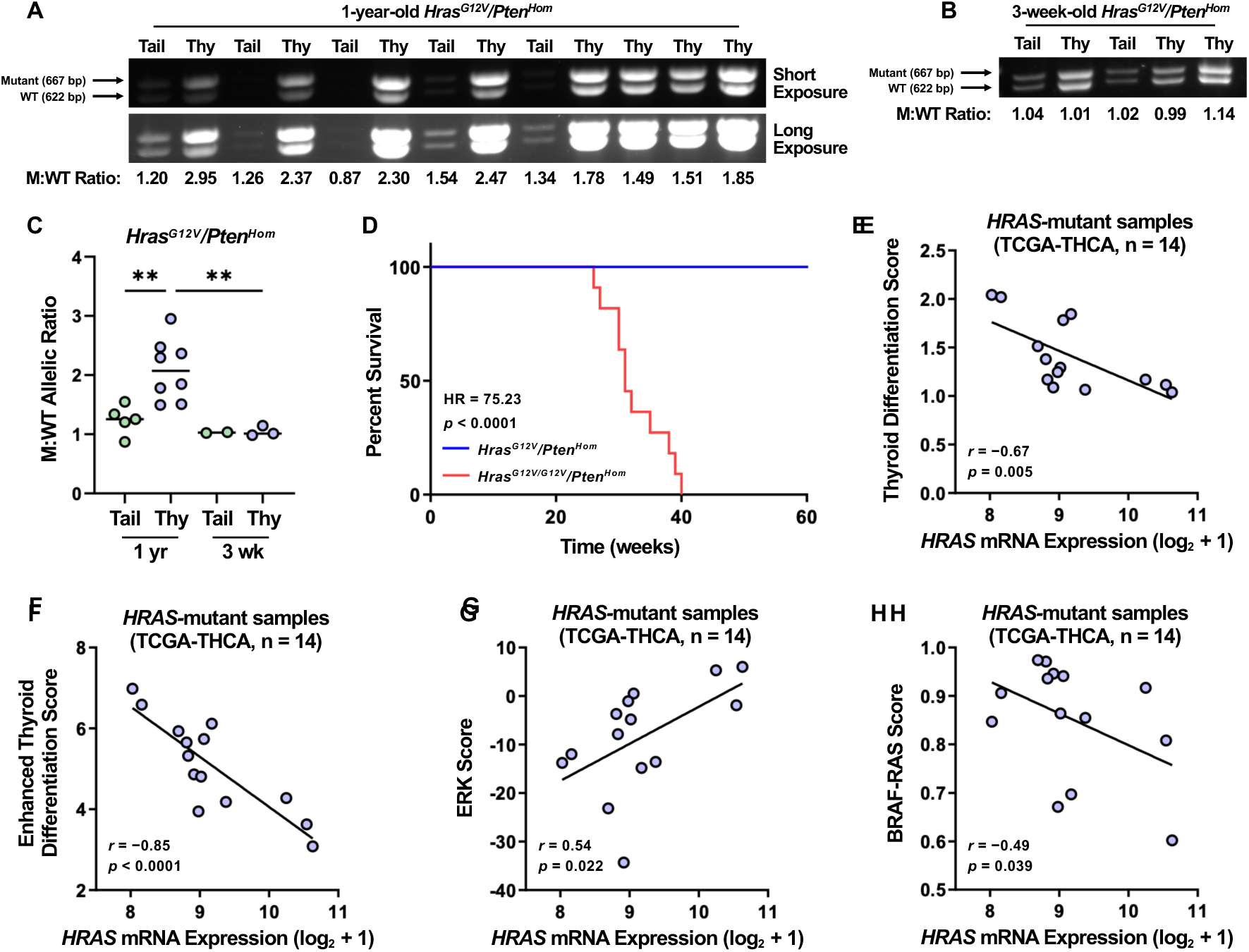
**(A-B)** PCR analysis of tail and thyroid genomic DNA from 1-year-old and 3-week-old *Hras^G12V^/Pten^Hom^* mice using primers amplifying both mutant (upper band; 667 bp) and wild-type (lower band; 622 bp) *Hras* alleles. The ratio of mutant to wild-type (M:WT) band densitometry is indicated. **(C)** Quantification of the mutant-to-wild-type allelic ratio in tail and thyroid DNA from 1-year-old and 3-week-old *Hras^G12V^/Pten^Hom^* mice. **(D)** Kaplan-Meier survival analysis of *Hras^G12V^/Pten^Hom^* (heterozygous *Hras^G12V^* knock-in) and *Hras^G12V/G12V^/Pten^Hom^* (homozygous *Hras^G12V^* knock-in) mice. Statistical significance was determined using the log-rank (Mantel-Cox) test. **(E-H)** Scatter plots of thyroid differentiation score, enhanced thyroid differentiation score, ERK score, and BRAF-RAS score versus *HRAS* mRNA expression in *HRAS*-mutant human thyroid cancer samples from the TCGA-THCA cohort. Pearson’s correlation coefficient (r) and associated *p*-values are indicated on each plot.

To determine whether *HRAS* dosage is similarly associated with malignant transformation in human thyroid cancers, we analyzed molecular data from *HRAS*-mutant PTC tumors (14/509) in the TCGA-THCA dataset. In contrast to the frequent allelic imbalance observed in our *Hras^G12V^*-driven animal model, only one *HRAS*-mutant human sample harbored increased variant allele frequency or copy-number gain. Given the rarity of genomic amplification in these samples, we evaluated *HRAS* dosage at a transcriptional level. Correlations between *HRAS* mRNA expression and established thyroid transcriptional scores associated with tumor differentiation and MAPK pathway activity, including thyroid differentiation (TDS), enhanced thyroid differentiation (eTDS), and ERK scores were assessed (30,31). Increased *HRAS* expression was associated with decreased levels of thyroid differentiation (TDS: r = -0.67, p = 0.005; eTDS: r = -0.85, p < 0.0001) in *HRAS*-mutant samples (Figure 4E, F). Further, *HRAS* expression was positively correlated with the ERK score of *HRAS-*mutant samples (r = 0.54, p = 0.022), suggesting that high *HRAS* expression may lead to increased MAPK output, presumably through increased expression of the mutant *HRAS* allele (Figure 4G). This is consistent with previous studies showing that increased expression of HRAS correlates with higher levels of pERK and pMEK (43). High *HRAS* expression was also associated with a decreased BRAF-RAS score (r = -0.49, p = 0.039), indicative of a more *BRAF^V600E^*-like transcriptional profile (Figure 4H). We then evaluated whether *KRAS* showed similar dose-dependent associations with TDS, eTDS, ERK, and BRAF-RAS scores in *KRAS*-mutant samples. Changes in these scores were not significantly correlated with *KRAS* expression, suggesting that *KRAS* dosage is not strongly associated with differentiation or MAPK output (Figure S6). However, these analyses are limited by the small number of *KRAS*-mutant tumors in the TCGA-THCA dataset (4/509). These associations suggest that *HRAS* dosage may contribute to a less differentiated, MAPK-active transcriptional state in human *HRAS*-mutant thyroid cancers, consistent with the dosage-dependent effects observed in our mouse model.

## Discussion

MAPK pathway activation, often through gain-of-function mutations in *RAS* family members, acts as a common driver of FTC and other thyroid tumors. However, while previous work has explored how broad MAPK activation cooperates with PI3K dysregulation to drive disease, the specific role of oncogenic *HRAS* in tumor initiation and progression has been understudied. To investigate how a common *HRAS* driver alteration cooperates with PI3K pathway activation to drive thyroid tumorigenesis, we developed a novel mouse model of FTC with thyroid-specific knock-in of *Hras^G12V^* and concurrent deletion of *Pten*. Critically, this model system, which recapitulates characteristics of human disease, allowed for the direct comparison of *Kras*- and *Hras*-mediated thyroid transformation. Our findings identified key molecular features of *Hras* and *Kras*-driven tumors that may mediate the observed variations in tumor latency between the *Hras^G12V^/Pten^Hom^* and *Kras^G12D^/Pten^Hom^* models.

As demonstrated in previous studies, germline expression of oncogenic *Hras* at physiological levels induces the development of hyperplastic lesions, squamous papillomas, and angiosarcomas (24). However, thyroid transformation was not observed in these mice or in mice with thyroid-specific expression of *Hras^G12V^*, even under chronic TSH stimulation, a potent mitogenic driver of thyroid growth (32). Indeed, it has been well reported that RAS-mediated transformation often requires cooperation with a second oncogene (44). Previous studies have similarly demonstrated that oncogenic *Kras* expression alone is insufficient to transform murine thyrocytes and that concurrent PI3K activation through *Pten* loss is required for complete transformation (20). Consistently, we found that thyrocyte-specific knock-in of *Hras^G12V^* combined with *Pten* loss induced robust thyroid transformation. This transformation was dependent on *Pten* dosage, with homozygous loss resulting in a higher incidence of malignant disease and a greater proportion of poorly differentiated tumors than mice with heterozygous loss. The resultant tumors recapitulated the histopathological features of human FTC and PDTC, while mice retained normal thyroid hormone biosynthesis, consistent with the observation that FTC patients rarely exhibit dysregulated thyroid hormone production (45–47). Together, these findings demonstrate that this model, which incorporates genetic alterations commonly found in human thyroid tumors, faithfully recapitulates key pathological and functional features of human disease. Interestingly, we also found that genetic background played a critical role in tumor development and malignancy. *Hras^G12V^/Pten^Hom^* mice on a pure 129SvJ background were more likely to develop malignant disease than their mixed-background counterparts, suggesting that there may be strain-specific genetic modifiers that impact penetrance of disease. Phenotypic differences between mouse strains have been widely reported within biological research, highlighting the importance of selecting proper model systems (48–51).

Previous studies have shown that different RAS isoforms exhibit distinct and often context-dependent transforming potential, demonstrating the relevance of evaluating RAS isoform function within specific tissue contexts (52–56). In the colon, expression of *KRAS^G12D^* induces hyperproliferation and increased tumorigenesis, whereas *NRAS^G12D^* does not (52). Conversely, in hematopoietic cells, oncogenic *NRAS* and *HRAS* induce more severe phenotypes and greater myeloproliferative disease compared with oncogenic *KRAS* (54,55). Similarly, *NRAS* has been shown to exhibit greater oncogenic potency than *KRAS* in melanocytes (56). Consistent with these findings, we identified differences in the oncogenic potency of *Hras^G12V^* and *Kras^G12D^* in the thyroid epithelium. While both mutations cooperated with homozygous *Pten* loss to induce FTC and PDTC, *Hras-*driven tumors displayed delayed tumor development compared with *Kras*-driven tumors. Despite the ability of *Kras^G12D^* to more rapidly transform thyrocytes, it remains unclear why *HRAS* mutations predominate in human thyroid tumors.

Having identified clear phenotypic differences induced by *Hras^G12V^* and *Kras^G12D^,* we sought to determine whether these phenotypes correlated with any molecular or biochemical differences. Previous studies have shown conflicting evidence regarding which RAS isoform more robustly activates the MAPK pathway, with these differences likely being dependent on cellular context (52,57–60). In the thyroid, we found that *Kras^G12D^* induces more robust ERK phosphorylation than *Hras^G12V^*, both during tumor initiation and following complete transformation, indicating that *Kras^G12D^* more potently and rapidly activates the MAPK pathway. Given that these studies were performed in the background of homozygous *Pten* loss, it is possible that constitutive PI3K pathway activation may influence the signaling dynamics of the two isoforms. Interestingly, *Kras^G12D^*-mutant thyroids exhibited significant downregulation of multiple MAPK negative regulators, including DUSP family phosphatases, during early tumorigenesis, suggesting a potential mechanism by which *Kras^G12D^* amplifies MAPK signaling by overcoming basal negative feedback. This early downregulation of DUSPs, which was not observed in *Hras-* mutant thyroids, may contribute to the rapid tumor formation observed in the *Kras*-driven model. While numerous studies have examined the basal signaling differences between oncogenic *Hras* and *Kras*, relatively few have investigated their differential sensitivities to MAPK pathway inhibition. One study evaluated the efficacy of MET inhibition in cells with ectopic expression of oncogenic HRAS and KRAS (61). However, given that MET signals upstream of RAS, the cells were expectedly resistant to MET inhibition. We therefore tested the sensitivities of oncogenic *Hras-* and *Kras-*driven cells to inhibition of downstream MAPK signaling. Interestingly, *Kras^G12D^* conferred reduced sensitivity to MEK inhibition compared with *Hras^G12V^*, possibly due to the higher basal MAPK and PI3K pathway activity induced by *Kras^G12D^*. Future studies should evaluate whether these trends are conserved across other cell types and with inhibition of other MAPK pathway proteins, including RAF and ERK, to better define the clinical relevance of these findings.

Copy number amplification is a common mechanism of oncogene activation, including for RAS family genes (62). Indeed, previous work demonstrated that amplification of *Hras^G12V^* is required to induce complete transformation in the skin (24,42). Consistent with these findings, we observed mutant *Hras* amplification in thyroid tumors from 1-year-old mice, but not in 3-week-old mice, indicating that this amplification is acquired during transformation. Interestingly, *Kras^G12D^/Pten^Hom^* mice retained allelic balance throughout tumor development, suggesting that while *Hras* may require increased allele dosage to transform thyrocytes, mutant *Kras* is sufficiently potent without additional amplification. This finding is distinct from other cancer types, where *KRAS^G12D^* amplification has been shown to promote tumor initiation, further highlighting the tissue-dependent properties of RAS isoforms (63,64). While *HRAS* copy-number gain was infrequent among PTC samples in the TCGA dataset, *HRAS* expression correlated with increased MAPK output and decreased differentiation. Interestingly, all samples in the TCGA dataset harbored *HRAS^Q61^* mutations, suggesting that the effect of *HRAS* dosage may not be unique to a specific mutant codon. As *RAS* alterations are more commonly associated with FTC, which are not included in the TCGA-THCA dataset, future studies should evaluate *RAS* allelic imbalance in large-scale human FTC cohorts (11,30). Together, our findings indicate that increased *HRAS* dosage, whether through genomic amplification or transcriptomic expression, plays a critical role in downstream signaling and tumor progression.

In this study, we present a murine model of FTC with thyroid-specific expression of oncogenic *Hras* and *Pten* loss. Although *HRAS^Q61^* mutations are more commonly observed in thyroid tumors, we utilized *Hras^G12V^* to enable direct comparison with the established *Kras^G12D^/Pten^Hom^* model and to assess the transforming properties of the corresponding G12 mutations across RAS isoforms (21,65). Here, we identified isoform-specific differences in oncogenic capacity and probed associated molecular differences that may underlie these phenotypes. Future studies should evaluate how specific codon mutations across all four RAS isoforms differentially activate the MAPK pathway and induce transformation in the thyroid gland.

## Acknowledgements

We would like to thank Dr. James Fagin at Memorial Sloan Kettering Cancer Center for sharing the *Hras*/*Pten* mice following ATF’s fellowship in the Fagin lab, and for the Fagin lab for insightful comments and critique during the initial stages of this project. We thank Samual Refetoff and the Refetoff laboratory for assistance with the TSH and T4 assays. **Grant support:** This work was supported in part through The University of Arkansas for Medical Sciences CTSA grant NID UL1TR000039; The National Institute of General Medical Sciences supported this work through the Center for Microbial Pathogenesis and Host Inflammatory Responses at the University of Arkansas for Medical Sciences COBRE Grant 1P20GM103625-02: American Thyroid Association/Thyca research grant (A. Franco); UAMS Envoys Seeds of Science Award (A. Franco) and by NIH grants (F32CA136178:A. Franco, R01CA214511:A. Franco, R01CA128943:A.DiCristofano). This content is soley the responsibility of the authors and does not necessarily represent the official views of the Nation Institutes of Health.

## References

1. Lim H, Devesa SS, Sosa JA, Check D, Kitahara CM. Trends in Thyroid Cancer Incidence and Mortality in the United States, 1974-2013. JAMA : the journal of the American Medical Association. 2017;317(13):1338–1348.

2. Davies L, Welch HG. Current thyroid cancer trends in the United States. JAMA Otolaryngol Head Neck Surg. 2014;140(4):317–322.

3. Rahib L, Smith BD, Aizenberg R, Rosenzweig AB, Fleshman JM, Matrisian LM. Projecting cancer incidence and deaths to 2030: the unexpected burden of thyroid, liver, and pancreas cancers in the United States. Cancer Res. 2014;74(11):2913–2921.

4. Hayat MJ, Howlader N, Reichman ME, Edwards BK. Cancer statistics, trends, and multiple primary cancer analyses from the Surveillance, Epidemiology, and End Results (SEER) Program. Oncologist. 2007;12(1):20–37.

5. Silver Karcioglu A, Slough C, Jordan SV, Brown TD, Kenzie JRM, Campbell DT, Dennis D, Kinua AG, Spaulding SL, Goli A, Bible K, Freeman JL, Newbold K, Shaha AR, Urken M, Zafereo M, Abdelhamid Ahmed AH, Randolph GW. Anaplastic Thyroid Carcinoma: A Contemporary Review of Challenges and Advances. Endocr Pract. 2026;32(1):118–126.

6. Ibrahimpasic T, Ghossein R, Shah JP, Ganly I. Poorly Differentiated Carcinoma of the Thyroid Gland: Current Status and Future Prospects. Thyroid : official journal of the American Thyroid Association. 2019;29(3):311–321.

7. Nikiforov YE, Nikiforova MN. Molecular genetics and diagnosis of thyroid cancer. Nat Rev Endocrinol. 2011;7(10):569–580.

8. Landa I, Ibrahimpasic T, Boucai L, Sinha R, Knauf JA, Shah RH, Dogan S, Ricarte-Filho JC, Krishnamoorthy GP, Xu B, Schultz N, Berger MF, Sander C, Taylor BS, Ghossein R, Ganly I, Fagin JA. Genomic and transcriptomic hallmarks of poorly differentiated and anaplastic thyroid cancers. The Journal of Clinical Investigation. 2016;126(3):1052– 1066.

9. Landa I, Pozdeyev N, Korch C, Marlow LA, Smallridge RC, Copland JA, Henderson YC, Lai SY, Clayman GL, Onoda N, Tan AC, Garcia-Rendueles MER, Knauf JA, Haugen BR, Fagin JA, Schweppe RE. Comprehensive Genetic Characterization of Human Thyroid Cancer Cell Lines: A Validated Panel for Preclinical Studies. Clinical Cancer Research. 2019;25(10):3141–3151.

10. Pozdeyev N, Gay LM, Sokol ES, Hartmaier R, Deaver KE, Davis S, French JD, Borre PV, LaBarbera DV, Tan A-C, Schweppe RE, Fishbein L, Ross JS, Haugen BR, Bowles DW. Genetic Analysis of 779 Advanced Differentiated and Anaplastic Thyroid Cancers. Clinical Cancer Research. 2018;24(13):3059–3068.

11. Nikiforov YE. Molecular analysis of thyroid tumors. Mod Pathol. 2011;24(S2):S34–S43.

12. Garcia-Rostan G, Costa AM, Pereira-Castro I, Salvatore G, Hernandez R, Hermsem MJ, Herrero A, Fusco A, Cameselle-Teijeiro J, Santoro M. Mutation of the PIK3CA gene in anaplastic thyroid cancer. Cancer Res. 2005;65(22):10199–10207.

13. Wu G, Mambo E, Guo Z, Hu S, Huang X, Gollin SM, Trink B, Ladenson PW, Sidransky D, Xing M. Uncommon mutation, but common amplifications, of the PIK3CA gene in thyroid tumors. The Journal of clinical endocrinology and metabolism. 2005;90(8):4688– 4693.

14. Ricarte-Filho JC, Ryder M, Chitale DA, Rivera M, Heguy A, Ladanyi M, Janakiraman M, Solit D, Knauf JA, Tuttle RM, Ghossein RA, Fagin JA. Mutational profile of advanced primary and metastatic radioactive iodine-refractory thyroid cancers reveals distinct pathogenetic roles for BRAF, PIK3CA, and AKT1. Cancer Res. 2009;69(11):4885–4893.

15. Hou P, Liu D, Shan Y, Hu S, Studeman K, Condouris S, Wang Y, Trink A, El-Naggar AK, Tallini G, Vasko V, Xing M. Genetic alterations and their relationship in the phosphatidylinositol 3-kinase/Akt pathway in thyroid cancer. Clin Cancer Res. 2007;13(4):1161–1170.

16. Liu Z, Hou P, Ji M, Guan H, Studeman K, Jensen K, Vasko V, El-Naggar AK, Xing M. Highly prevalent genetic alterations in receptor tyrosine kinases and phosphatidylinositol 3-kinase/akt and mitogen-activated protein kinase pathways in anaplastic and follicular thyroid cancers. The Journal of clinical endocrinology and metabolism. 2008;93(8):3106–3116.

17. Saji M, Ringel MD. The PI3K-Akt-mTOR pathway in initiation and progression of thyroid tumors. Mol Cell Endocrinol. 2010;321(1):20–28.

18. Baran JA, Tsai SD, Isaza A, Brodeur GM, MacFarland SP, Zelley K, Adams DM, Franco AT, Bauer AJ. The Clinical Spectrum of PTEN Hamartoma Tumor Syndrome: Exploring the Value of Thyroid Surveillance. Horm Res Paediatr. 2020;93(11-12):634–642.

19. Jolly LA, Novitskiy S, Owens P, Massoll N, Cheng N, Fang W, Moses HL, Franco AT. Fibroblast-Mediated Collagen Remodeling Within the Tumor Microenvironment Facilitates Progression of Thyroid Cancers Driven by BrafV600E and Pten Loss. Cancer Res. 2016;76(7):1804–1813.

20. Miller KA, Yeager N, Baker K, Liao XH, Refetoff S, Di Cristofano A. Oncogenic Kras requires simultaneous PI3K signaling to induce ERK activation and transform thyroid epithelial cells in vivo. Cancer Res. 2009;69(8):3689–3694.

21. Volante M, Rapa I, Gandhi M, Bussolati G, Giachino D, Papotti M, Nikiforov YE. RAS Mutations Are the Predominant Molecular Alteration in Poorly Differentiated Thyroid Carcinomas and Bear Prognostic Impact. Journal of Clinical Endocrinology & Metabolism. 2009;94(12):4735–4741.

22. Boucai L, Seshan V, Williams M, Knauf JA, Saqcena M, Ghossein RA, Fagin JA. Characterization of Subtypes of BRAF-Mutant Papillary Thyroid Cancer Defined by Their Thyroid Differentiation Score. The Journal of clinical endocrinology and metabolism. 2022;107(4):1030–1039.

23. Na KJ, Choi H. Immune landscape of papillary thyroid cancer and immunotherapeutic implications. Endocr Relat Cancer. 2018;25(5):523–531.

24. Chen X, Mitsutake N, LaPerle K, Akeno N, Zanzonico P, Longo VA, Mitsutake S, Kimura ET, Geiger H, Santos E, Wendel HG, Franco A, Knauf JA, Fagin JA. Endogenous expression of Hras(G12V) induces developmental defects and neoplasms with copy number imbalances of the oncogene. Proceedings of the National Academy of Sciences of the United States of America. 2009;106(19):7979–7984.

25. Kusakabe T, Kawaguchi A, Kawaguchi R, Feigenbaum L, Kimura S. Thyrocyte-specific expression of Cre recombinase in transgenic mice. Genesis. 2004;39(3):212–216.

26. Trotman LC, Niki M, Dotan ZA, Koutcher JA, Di Cristofano A, Xiao A, Khoo AS, Roy-Burman P, Greenberg NM, Van Dyke T, Cordon-Cardo C, Pandolfi PP. Pten dose dictates cancer progression in the prostate. PLoS biology. 2003;1(3):E59.

27. Pohlenz J, Maqueem A, Cua K, Weiss RE, Van Sande J, Refetoff S. Improved radioimmunoassay for measurement of mouse thyrotropin in serum: strain differences in thyrotropin concentration and thyrotroph sensitivity to thyroid hormone. Thyroid : official journal of the American Thyroid Association. 1999;9(12):1265–1271.

28. Dima M, Miller KA, Antico-Arciuch VG, Di Cristofano A. Establishment and characterization of cell lines from a novel mouse model of poorly differentiated thyroid carcinoma: powerful tools for basic and preclinical research. Thyroid : official journal of the American Thyroid Association. 2011;21(9):1001–1007.

29. Cerami E, Gao J, Dogrusoz U, Gross BE, Sumer SO, Aksoy BA, Jacobsen A, Byrne CJ, Heuer ML, Larsson E, Antipin Y, Reva B, Goldberg AP, Sander C, Schultz N. The cBio cancer genomics portal: an open platform for exploring multidimensional cancer genomics data. Cancer discovery. 2012;2(5):401–404.

30. 30. Integrated genomic characterization of papillary thyroid carcinoma. Cell. 2014;159(3):676–690.

31. Dunn LA, Sherman EJ, Baxi SS, Tchekmedyian V, Grewal RK, Larson SM, Pentlow KS, Haque S, Tuttle RM, Sabra MM, Fish S, Boucai L, Walters J, Ghossein RA, Seshan VE, Ni A, Li D, Knauf JA, Pfister DG, Fagin JA, Ho AL. Vemurafenib Redifferentiation of BRAF Mutant, RAI-Refractory Thyroid Cancers. The Journal of clinical endocrinology and metabolism. 2019;104(5):1417–1428.

32. Franco AT, Malaguarnera R, Refetoff S, Liao XH, Lundsmith E, Kimura S, Pritchard C, Marais R, Davies TF, Weinstein LS, Chen M, Rosen N, Ghossein R, Knauf JA, Fagin JA. Thyrotrophin receptor signaling dependence of Braf-induced thyroid tumor initiation in mice. Proceedings of the National Academy of Sciences of the United States of America. 2011;108(4):1615–1620.

33. Jhiang SM, Sagartz JE, Tong Q, Parker-Thornburg J, Capen CC, Cho JY, Xing S, Ledent C. Targeted expression of the ret/PTC1 oncogene induces papillary thyroid carcinomas. Endocrinology. 1996;137(1):375–378.

34. Knauf JA, Ma X, Smith EP, Zhang L, Mitsutake N, Liao XH, Refetoff S, Nikiforov YE, Fagin JA. Targeted expression of BRAFV600E in thyroid cells of transgenic mice results in papillary thyroid cancers that undergo dedifferentiation. Cancer Res. 2005;65(10):4238–4245.

35. Li W, Reeb AN, Sewell WA, Elhomsy G, Lin RY. Phenotypic characterization of metastatic anaplastic thyroid cancer stem cells. PloS one. 2013;8(5):e65095.

36. Balko JM, Schwarz LJ, Bhola NE, Kurupi R, Owens P, Miller TW, Gómez H, Cook RS, Arteaga CL. Activation of MAPK Pathways due to DUSP4 Loss Promotes Cancer Stem Cell-like Phenotypes in Basal-like Breast Cancer. Cancer Research. 2013;73(20):6346– 6358.

37. Bermudez O, Pages G, Gimond C. The dual-specificity MAP kinase phosphatases: critical roles in development and cancer. American journal of physiology Cell physiology. 2010;299(2):C189–202.

38. Caunt CJ, Keyse SM. Dual-specificity MAP kinase phosphatases (MKPs). FEBS Journal. 2013;280(2):489–504.

39. Keyse S. Dual-specificity MAP kinase phosphatases (MKPs) and cancer. Cancer Metastasis Rev. 2008;27(2):253–261.

40. Schmid CA, Robinson MD, Scheifinger NA, Müller S, Cogliatti S, Tzankov A, Müller A. DUSP4 deficiency caused by promoter hypermethylation drives JNK signaling and tumor cell survival in diffuse large B cell lymphoma. The Journal of Experimental Medicine. 2015;212(5):775–792.

41. Sieben NLG, Oosting J, Flanagan AM, Prat J, Roemen GMJM, Kolkman-Uljee SM, van Eijk R, Cornelisse CJ, Fleuren GJ, van Engeland M. Differential Gene Expression in Ovarian Tumors Reveals Dusp 4 and Serpina 5 As Key Regulators for Benign Behavior of Serous Borderline Tumors. Journal of Clinical Oncology. 2005;23(29):7257–7264.

42. Chen X, Makarewicz JM, Knauf JA, Johnson LK, Fagin JA. Transformation by Hras(G12V) is consistently associated with mutant allele copy gains and is reversed by farnesyl transferase inhibition. Oncogene. 2014;33(47):5442–5449.

43. Garcia-Rendueles ME, Ricarte-Filho JC, Untch BR, Landa I, Knauf JA, Voza F, Smith VE, Ganly I, Taylor BS, Persaud Y, Oler G, Fang Y, Jhanwar SC, Viale A, Heguy A, Huberman KH, Giancotti F, Ghossein R, Fagin JA. NF2 Loss Promotes Oncogenic RAS-Induced Thyroid Cancers via YAP-Dependent Transactivation of RAS Proteins and Sensitizes Them to MEK Inhibition. Cancer discovery. 2015;5(11):1178–1193.

44. Land H, Parada LF, Weinberg RA. Tumorigenic conversion of primary embryo fibroblasts requires at least two cooperating oncogenes. Nature. 1983;304(5927):596–602.

45. Lee EK, Chung KW, Min HS, Kim TS, Kim TH, Ryu JS, Jung YS, Kim SK, Lee YJ. Preoperative serum thyroglobulin as a useful predictive marker to differentiate follicular thyroid cancer from benign nodules in indeterminate nodules. Journal of Korean medical science. 2012;27(9):1014–1018.

46. Lee SH, Baek JS, Lee JY, Lim JA, Cho SY, Lee TH, Ku YH, Kim HI, Kim MJ. Predictive factors of malignancy in thyroid nodules with a cytological diagnosis of follicular neoplasm. Endocrine pathology. 2013;24(4):177–183.

47. Rinaldi S, Plummer M, Biessy C, Tsilidis KK, Ostergaard JN, Overvad K, Tjonneland A, Halkjaer J, Boutron-Ruault MC, Clavel-Chapelon F, Dossus L, Kaaks R, Lukanova A, Boeing H, Trichopoulou A, Lagiou P, Trichopoulos D, Palli D, Agnoli C, Tumino R, Vineis P, Panico S, Bueno-de-Mesquita HB, Peeters PH, Weiderpass E, Lund E, Quiros JR, Agudo A, Molina E, Larranaga N, Navarro C, Ardanaz E, Manjer J, Almquist M, Sandstrom M, Hennings J, Khaw KT, Schmidt J, Travis RC, Byrnes G, Scalbert A, Romieu I, Gunter M, Riboli E, Franceschi S. Thyroid-stimulating hormone, thyroglobulin, and thyroid hormones and risk of differentiated thyroid carcinoma: the EPIC study. Journal of the National Cancer Institute. 2014;106(6):dju097.

48. Bufi R, Korstanje R. The impact of genetic background on mouse models of kidney disease. Kidney Int. 2022;102(1):38–44.

49. Gurram RK, Wei D, Yu Q, Butcher MJ, Chen X, Cui K, Hu G, Zheng M, Zhu X, Oh J, Sun B, Urban JF, Jr., Zhao K, Leonard WJ, Zhu J. Crosstalk between ILC2s and Th2 cells varies among mouse models. Cell Rep. 2023;42(2):112073.

50. Norris AM, Fierman KE, Campbell J, Pitale R, Shahraj M, Kopinke D. Studying intramuscular fat deposition and muscle regeneration: insights from a comparative analysis of mouse strains, injury models, and sex differences. Skelet Muscle. 2024;14(1):12.

51. Olguin V, Duran A, Las Heras M, Rubilar JC, Cubillos FA, Olguin P, Klein AD. Genetic Background Matters: Population-Based Studies in Model Organisms for Translational Research. Int J Mol Sci. 2022;23(14).

52. Haigis KM, Kendall KR, Wang Y, Cheung A, Haigis MC, Glickman JN, Niwa-Kawakita M, Sweet-Cordero A, Sebolt-Leopold J, Shannon KM, Settleman J, Giovannini M, Jacks T. Differential effects of oncogenic K-Ras and N-Ras on proliferation, differentiation and tumor progression in the colon. Nat Genet. 2008;40(5):600–608.

53. Fotiadou PP, Takahashi C, Rajabi HN, Ewen ME. Wild-type NRas and KRas perform distinct functions during transformation. Mol Cell Biol. 2007;27(19):6742–6755.

54. Parikh C, Subrahmanyam R, Ren R. Oncogenic NRAS, KRAS, and HRAS exhibit different leukemogenic potentials in mice. Cancer Res. 2007;67(15):7139–7146.

55. Xu J, Haigis KM, Firestone AJ, McNerney ME, Li Q, Davis E, Chen SC, Nakitandwe J, Downing J, Jacks T, Le Beau MM, Shannon K. Dominant role of oncogene dosage and absence of tumor suppressor activity in Nras-driven hematopoietic transformation. Cancer discovery. 2013;3(9):993–1001.

56. Whitwam T, Vanbrocklin MW, Russo ME, Haak PT, Bilgili D, Resau JH, Koo HM, Holmen SL. Differential oncogenic potential of activated RAS isoforms in melanocytes. Oncogene. 2007;26(31):4563–4570.

57. Voice JK, Klemke RL, Le A, Jackson JH. Four human ras homologs differ in their abilities to activate Raf-1, induce transformation, and stimulate cell motility. J Biol Chem. 1999;274(24):17164–17170.

58. Rosseland CM, Wierod L, Flinder LI, Oksvold MP, Skarpen E, Huitfeldt HS. Distinct functions of H-Ras and K-Ras in proliferation and survival of primary hepatocytes due to selective activation of ERK and PI3K. J Cell Physiol. 2008;215(3):818–826.

59. Cespedes MV, Sancho FJ, Guerrero S, Parreno M, Casanova I, Pavon MA, Marcuello E, Trias M, Cascante M, Capella G, Mangues R. K-ras Asp12 mutant neither interacts with Raf, nor signals through Erk and is less tumorigenic than K-ras Val12. Carcinogenesis. 2006;27(11):2190–2200.

60. Yan J, Roy S, Apolloni A, Lane A, Hancock JF. Ras isoforms vary in their ability to activate Raf-1 and phosphoinositide 3-kinase. J Biol Chem. 1998;273(37):24052–24056.

61. Leiser D, Medova M, Mikami K, Nisa L, Stroka D, Blaukat A, Bladt F, Aebersold DM, Zimmer Y. KRAS and HRAS mutations confer resistance to MET targeting in preclinical models of MET-expressing tumor cells. Mol Oncol. 2015;9(7):1434–1446.

62. Bagci O, Kurtgoz S. Amplification of Cellular Oncogenes in Solid Tumors. N Am J Med Sci. 2015;7(8):341–346.

63. Mueller S, Engleitner T, Maresch R, Zukowska M, Lange S, Kaltenbacher T, Konukiewitz B, Ollinger R, Zwiebel M, Strong A, Yen HY, Banerjee R, Louzada S, Fu B, Seidler B, Gotzfried J, Schuck K, Hassan Z, Arbeiter A, Schonhuber N, Klein S, Veltkamp C, Friedrich M, Rad L, Barenboim M, Ziegenhain C, Hess J, Dovey OM, Eser S, Parekh S, Constantino-Casas F, de la Rosa J, Sierra MI, Fraga M, Mayerle J, Kloppel G, Cadinanos J, Liu P, Vassiliou G, Weichert W, Steiger K, Enard W, Schmid RM, Yang F, Unger K, Schneider G, Varela I, Bradley A, Saur D, Rad R. Evolutionary routes and KRAS dosage define pancreatic cancer phenotypes. Nature. 2018;554(7690):62–68.

64. Najumudeen AK, Fey SK, Millett LM, Ford CA, Gilroy K, Gunduz N, Ridgway RA, Anderson E, Strathdee D, Clark W, Nixon C, Morton JP, Campbell AD, Sansom OJ. KRAS allelic imbalance drives tumour initiation yet suppresses metastasis in colorectal cancer in vivo. Nat Commun. 2024;15(1):100.

65. Nikiforov YE. Molecular Diagnostics of Thyroid Tumors. Archives of Pathology & Laboratory Medicine. 2011;135(5):569–577.

