## Supplemental Figures for "*Hras^G12V^* induces follicular thyroid cancer with attenuated MAPK activation and increased latency compared to *Kras^G12D^*"

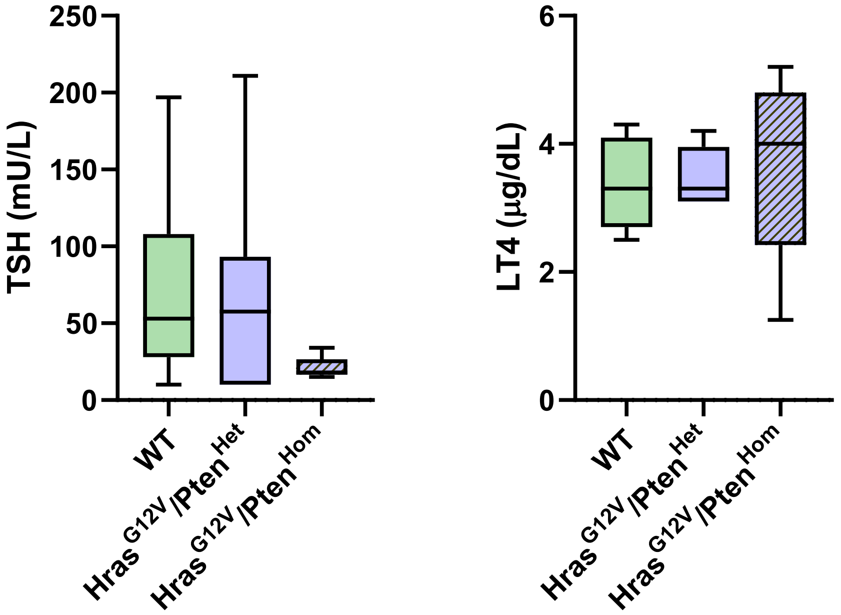


**Figure S1.** Levels of thyroid stimulating hormone (TSH) and levothyroxine (LT4) in serum from wild-type (n = 38), *Hras*$^{G12V}$*/Pten*$^{Het}$ (n = 18), and *Hras*$^{G12V}$*/Pten*$^{Hom}$ (n = 5) mice. There was no statistical significance between any groups as measured by one-way ANOVA.

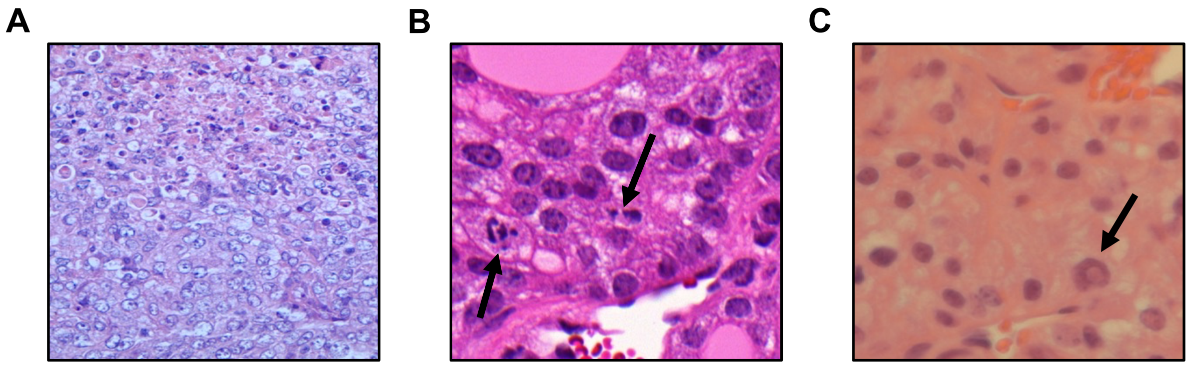


**Figure S2.** Additional images of H&E-stained thyroid tissue from *Hras*$^{G12V}$/*Pten*$^{Hom}$ mice. **(A)** 40X image showing follicular cells with vesicular nuclei with central necrosis. **(B)** 100X image showing follicular cells with atypical mitoses (arrows). **(C)** 100X image showing a follicular cell with nuclear psuedoinclusion (arrow).

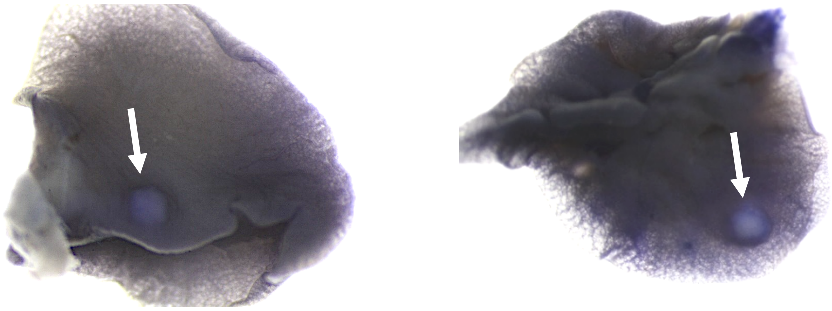

**Figure S3.** Representative images of lungs from *Hras*$^{G12V}$/*Pten*$^{Hom}$ mice harboring macrometastases (arrows).

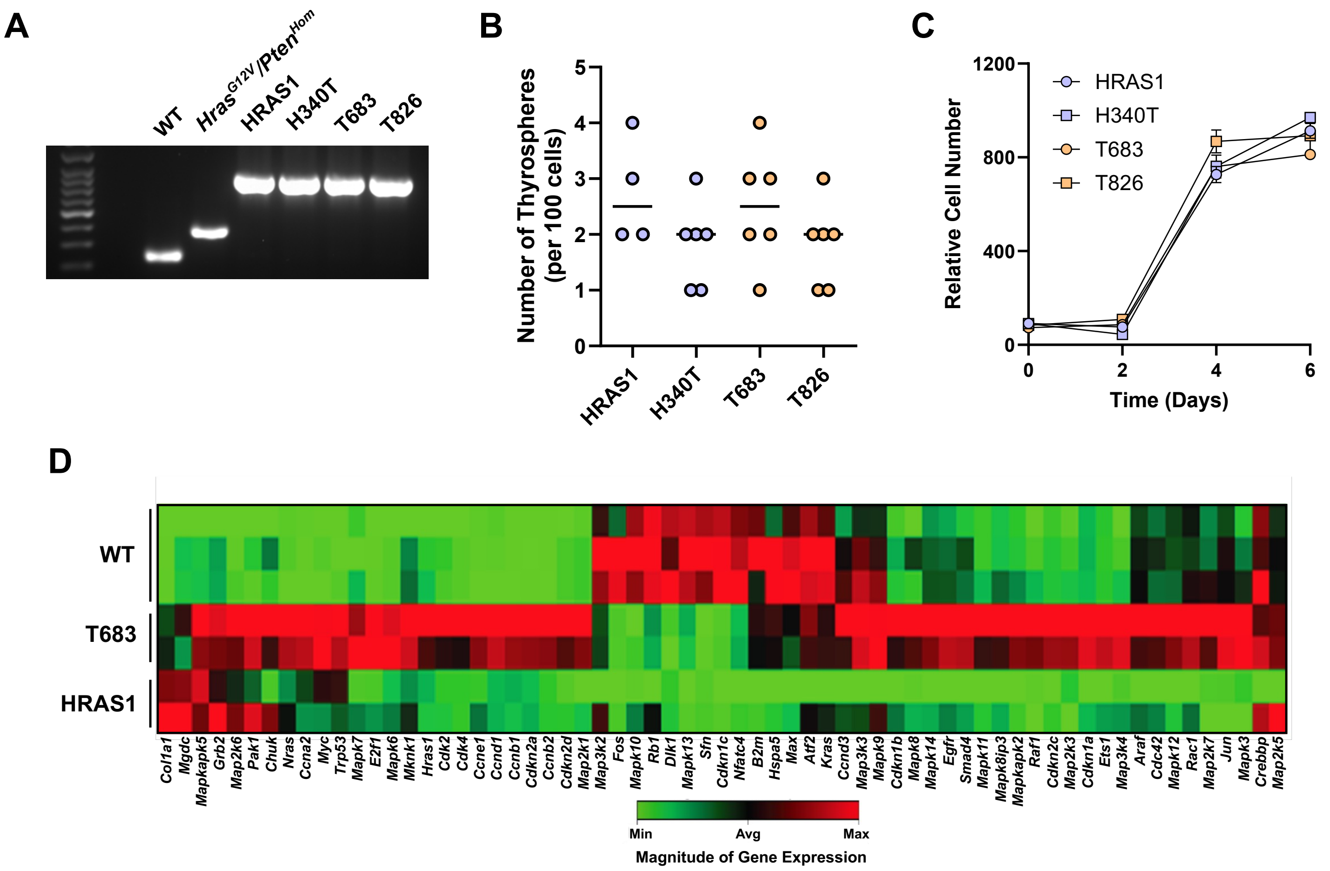


**Figure S4. (A)** PCR gel showing complete recombination of *Pten* in HRAS1, H340T, T683, and T826 cell lines. Tail DNA from wild-type and *$Hras^{G12V}/Pten^{Hom}$* mice were run as controls. **(B)** Number of thyrospheres formed per 100 cells plated from HRAS1, H340T, T683, and T826 cell lines. There was no statistically significant difference between cell lines as measured by one-way ANOVA. **(C)** Growth curves of HRAS1, H340T, T683, and T826 cell lines. Quantification was performed across ten replicates per cell line per day. **(D)** RT-qPCR array measuring the expression of 84 genes involved in the MAPK signaling pathway in wild-type thyroids (n = 3), T683 (n = 2), and HRAS1 (n = 2).

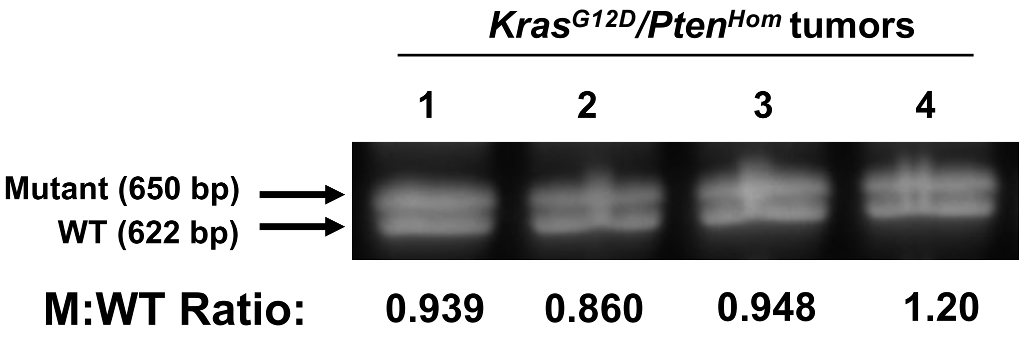


**Figure S5.** PCR analysis of thyroid genomic DNA from *Kras*$^{G12D}$/*Pten*$^{Hom}$ mice using primers amplifying both mutant (upper band; 650 bp) and wild-type (lower band; 622 bp) *Kras* alleles. The ratio of mutant to wild-type (M:WT) band densitometry is indicated below the gel.

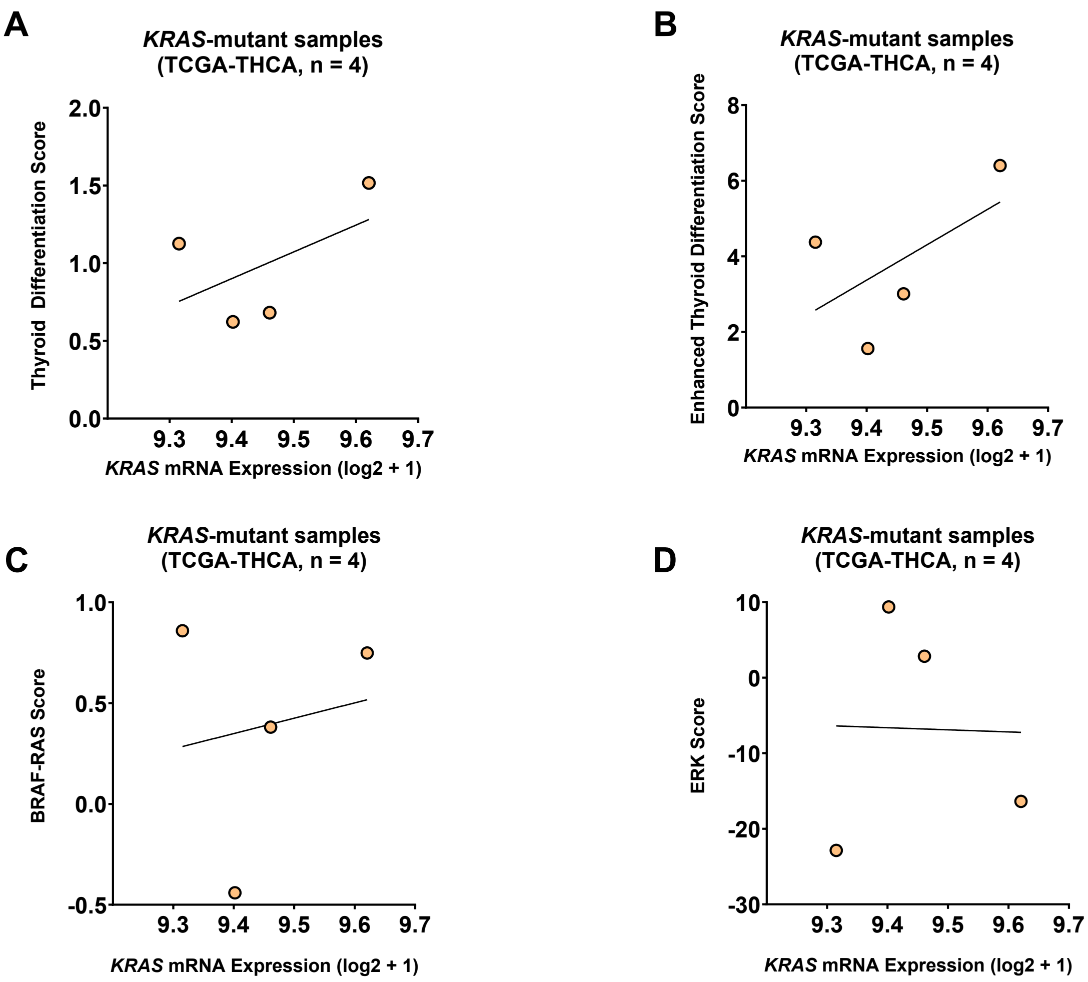


**Figure S6.** Scatter plots of **(A)** thyroid differentiation score, **(B)** enhanced thyroid differentiation score, **(C)** BRAF-RAS score, and **(D)** ERK score versus *KRAS* mRNA expression in *KRAS*-mutant human thyroid cancer samples from the TCGA-THCA cohort. No statistically significant correlation was found between *KRAS* mRNA expression and any molecular score.
